# Phylogenomics supports the reinstatement of *Sinia* (Ochnaceae)

**DOI:** 10.64898/2026.08.24.746610

**Authors:** Tian-Wen Xiao, Xue-Jun Ge

## Abstract

*Sinia rhodoleuca*, the sole species of the monotypic genus *Sinia* (Ochnaceae), was previously transferred to *Sauvagesia* based mainly on morphological similarities. However, its phylogenetic position has remained unresolved because molecular data for the species were unavailable. Here, we generated genomic data for *Sinia rhodoleuca* and reconstructed its phylogenetic position within Sauvagesieae. Our phylogenomic analyses consistently recovered *Sinia rhodoleuca* as sister to *Indosinia*, whereas the Neotropical *Sauvagesia* formed a distantly related lineage, rendering *Sauvagesia* broadly circumscribed non-monophyletic. Comparative morphological evidence further supports the close relationship between *Sinia* and *Indosinia*, particularly in their closely parallel secondary veins, lacerate stipules, and prominent petaloid staminodes, while differences in floral characters support their recognition as distinct genera. We therefore reinstate *Sinia* as a distinct genus and provide a revised taxonomic treatment of *Sinia rhodoleuca*. Our study demonstrates how phylogenomic evidence can resolve long-standing taxonomic uncertainties and reveal evolutionary relationships obscured by morphological similarity.

**Main conclusion:** Phylogenomic evidence supports the reinstatement of *Sinia* as a distinct genus and reveals its sister relationship with *Indosinia*, refining generic boundaries in Sauvagesieae.

## Introduction

Accurate inference of evolutionary relationships is fundamental to understanding plant diversity and establishing classifications that reflect evolutionary history. Phylogenomic approaches have substantially improved the resolution of relationships across angiosperms, particularly in lineages where morphological similarity and homoplasy complicate generic delimitation (Schneider et al. 2014; Song et al. 2020; Schneider et al. 2021b; Soares et al. 2021). In such groups, integrating genome-scale phylogenetic evidence with morphological data can help distinguish shared evolutionary history from independently evolved similarities and provide a more robust basis for taxonomic classification. The family Ochnaceae provides an example of this challenge. Within Ochnaceae, tribe Sauvagesieae comprises a morphologically diverse assemblage of tropical and subtropical lineages whose generic boundaries and intergeneric relationships have undergone considerable revision (Amaral 1991, 2006). Recent molecular and phylogenomic studies have substantially improved our understanding of relationships within the tribe, but several lineages remain poorly sampled or have unresolved taxonomic positions (Schneider et al. 2014; Schneider et al. 2021b; Reinales et al. 2026).

*Sinia* Diels is a monotypic genus of Sauvagesieae, originally described from southern China on the basis of *Sinia rhodoleuca* Diels (Diels 1930). The genus is characterized by closely parallel secondary veins, glandular marginal teeth, lacerate stipules, and a distinctive arrangement of petaloid staminodes. However, some of these characters are shared with other members of Sauvagesieae, and the taxonomic position of *Sinia* has consequently remained controversial (Amaral 2006; Kubitzki 2014; Schneider et al. 2021b). Based on similarities in floral and seed morphology, Amaral (2006) transferred *Sinia rhodoleuca* to *Sauvagesia*, thereby treating *Sinia* as a synonym of *Sauvagesia*. This treatment placed the species within a broadly circumscribed *Sauvagesia* that included morphologically diverse Neotropical and Asian lineages. Subsequent molecular studies, however, have shown that several taxa historically included in *Sauvagesia* are not closely related to its Neotropical species, suggesting that some morphological similarities used to delimit the broadly circumscribed genus may be homoplastic (Schneider et al. 2021b).

Despite these advances, the phylogenetic position of *Sinia* has remained unresolved. Schneider et al. (2014), Schneider et al. (2021b) and Reinales et al. (2026) substantially revised the phylogenetic framework of Sauvagesieae and demonstrated that several taxa formerly included in *Sauvagesia* represent independent evolutionary lineages. In particular, their results supported the recognition of *Indosinia, Indovethia*, and *Neckia* as distinct genera and revealed substantial discordance between morphology-based classifications and molecular relationships. However, *Sinia* was not included in the molecular analyses of Schneider et al. (2021b) because suitable material was unavailable, leaving its relationships to *Sauvagesia* and other Asian genera unresolved. This unresolved position is of particular taxonomic interest because the morphological characters that led to the inclusion of *Sinia* in *Sauvagesia* also occur, in different combinations, among other genera of Sauvagesieae (Kubitzki 2014; Schneider et al. 2021b). Resolving the position of *Sinia* is therefore important not only for determining its appropriate generic circumscription and assessing whether its similarities to *Sauvagesia* reflect common ancestry or homoplasy.

In this study, we generated genome-scale data using next generation sequencing (NGS) for *Sinia rhodoleuca* (≡*Sauvagesia rhodoleuca*) and incorporated these data into a phylogenomic framework of Ochnaceae to resolve its previously uncertain systematic position. We further examined key morphological characters, particularly leaf venation, stipules, staminodes, and anther dehiscence, to assess whether the inferred phylogenetic relationship is supported by morphology. Based on the combined phylogenomic and morphological evidence, we evaluate the generic status of *Sinia* and its relationship to *Indosinia* and other members of Sauvagesieae. Our results support the recognition of *Sinia* as a distinct genus and refine the circumscription of *Sauvagesia*. More broadly, our study highlights the value of integrating phylogenomics and comparative morphology to resolve taxonomic uncertainty in morphologically complex plant lineages.

## Material and methods

### Plant sampling and sequencing

Leaf material of *Sinia rhodoleuca* was collected from Heishiding Provincial Nature Reserve in Fengkai city, Guangdong province, China, and immediately silica-dried. Voucher specimen (gexj230011) was deposited in the herbarium of the South China Botanical Garden, Chinese Academy of Sciences (IBSC).

Genomic DNA was extracted using the cetyltrimethylammonium bromide (CTAB) method (Doyle and Doyle 1987). A paired-end library (2 × 150 bp) was prepared using the TruSeq Nano DNA HT Sample Preparation Kit (Illumina, San Diego, CA, USA). Sequencing was performed on an MGI DNBSEQ-T7 platform at Grand Omics Co., Ltd., Wuhan, China.

### Nuclear gene assembly

Raw reads were quality-controlled using fastp v0.23.3 (Chen et al. 2018) with default parameters. Two nuclear datasets developed by Schneider et al. (2021b) were used to generate target files for HybPiper v2.1.1 (Johnson et al. 2016): the FAM dataset, which maximizes taxon sampling across Ochnaceae, and the SLT dataset, which includes only species from three tribes, Sauvagesieae, Luxemburgieae, and Testuleeae. Sequencing reads for 312 samples of Ochnaceae and outgroups from Schneider et al. (2021b) were downloaded from National Center for Biotechnology Information (NCBI) (Table S1). Clean reads from the 312 samples and *Sinia rhodoleuca* were assembled into single-copy nuclear genes (SCNs) using HybPiper v2.1.1 with default settings.

### Plastome assembly and annotation

Plastome of *Sinia rhodoleuca* was assembled using GetOrganelle v1.7.5.3 (Jin et al. 2020) and annotated using GeSeq (Tillich et al. 2017). Start and stop codons of protein-coding genes were manually checked and adjusted using Geneious v9.1.3 (Kearse et al. 2012). Circular map of plastome was visualized using PlastidHub (Zhang et al. 2025).

### Phylogenetic analyses

SCN sequences shorter than 100 bp were removed using SeqKit v2.3.0 (Shen et al. 2024). SCNs were aligned using MAFFT v7.508 (Katoh and Standley 2013) with default settings. Poorly aligned regions were removed using trimAl v1.4.1 (Capella-Gutiérrez et al. 2009) with the “automated1” option. Potential alignment errors were identified and removed using TAPER v1.0.0 (Zhang et al. 2021). For concatenation analyses, the processed SCN alignments were concatenated using AMAS v1.0 (Borowiec 2016), and maximum likelihood (ML) trees were inferred using RAxML v8.2.11 (Stamatakis 2014) under the GTRGAMMA model with 1,000 bootstrap replicates. For coalescent-based analyses, gene trees were inferred from the processed alignments using RAxML v8.2.11 under the GTRGAMMA model, with 200 bootstrap replicates. To minimize the effects of low support and/or long terminal branches in gene trees, weighted ASTRAL v1.15.2.3 (Zhang and Mirarab 2022) was used to infer the species tree.

For the plastid phylogenetic analysis, plastid DNA sequences of Ochnaceae were obtained from Schneider et al. (2021a). The plastid sequence of *Sinia rhodoleuca* and the reference dataset were aligned using MAFFT v7.508, and an ML tree was inferred using RAxML v8.2.11 under the GTRGAMMA model with 1,000 bootstrap replicates.

### Morphological assessment

Herbarium specimens and specimen images were examined digitally from the following herbaria: IBSC, K, M, S, BM, E, SBT, GDC, US, BR, PH, MA, and P. Additional specimen records and digital images were examined through online databases, including JSTOR Global Plants (https://plants.jstor.org/), the Chinese Virtual Herbarium (CVH, https://www.cvh.ac.cn), and the Plant Photo Bank of China (PPBC, https://ppbc.iplant.cn/). Morphological characters of *Sinia* and related genera were assessed based on published taxonomic treatments and descriptions (Amaral 2006; Zhang and Amaral 2007; Kubitzki 2014; Schneider et al. 2021b; Nguyen et al. 2025), with particular emphasis on leaf, floral, and seed characters. Particular attention was given to characters relevant to generic delimitation within Sauvagesieae, including stipule morphology, leaf venation, staminode arrangement, anther dehiscence, ovary structure, placentation, and seed morphology.

## Results

### Assembly of single-copy nuclear genes and plastomes

The newly sequenced sample of *Sinia rhodoleuca* yielded approximately 245.99 million reads, corresponding to 36.90 Gb of raw data. After quality control, 36.73 Gb of clean data were retained. A total of 46 and 63 SCNs were recovered for the FAM and SLT datasets, respectively.

The newly assembled plastome of *Sinia rhodoleuca* (GenBank accession number: PX933086) was 157,300 bp in length and exhibited a typical quadripartite structure consisting of a large single-copy region (LSC), a small single-copy region (SSC), and two inverted repeats (IRs) (Fig. S1).

### Phylogenetic position of *Sinia rhodoleuca*

Across both nuclear datasets and the different phylogenetic inference approaches, *Sinia rhodoleuca* was consistently recovered as sister to *Indosinia involucrata*, with maximal support (bootstrap = 100% and local posterior probability = 1; Fig. 1). The *Sinia rhodoleuca* + *Indosinia* clade was clearly separated from the sampled species of *Sauvagesia*. The latter formed two clades associated with *Adenarake muriculata* and *Tyleria*, respectively (Fig. 1). Thus, *Sinia rhodoleuca* was not closely related to the Neotropical *Sauvagesia* lineage.

**Figure 1.**
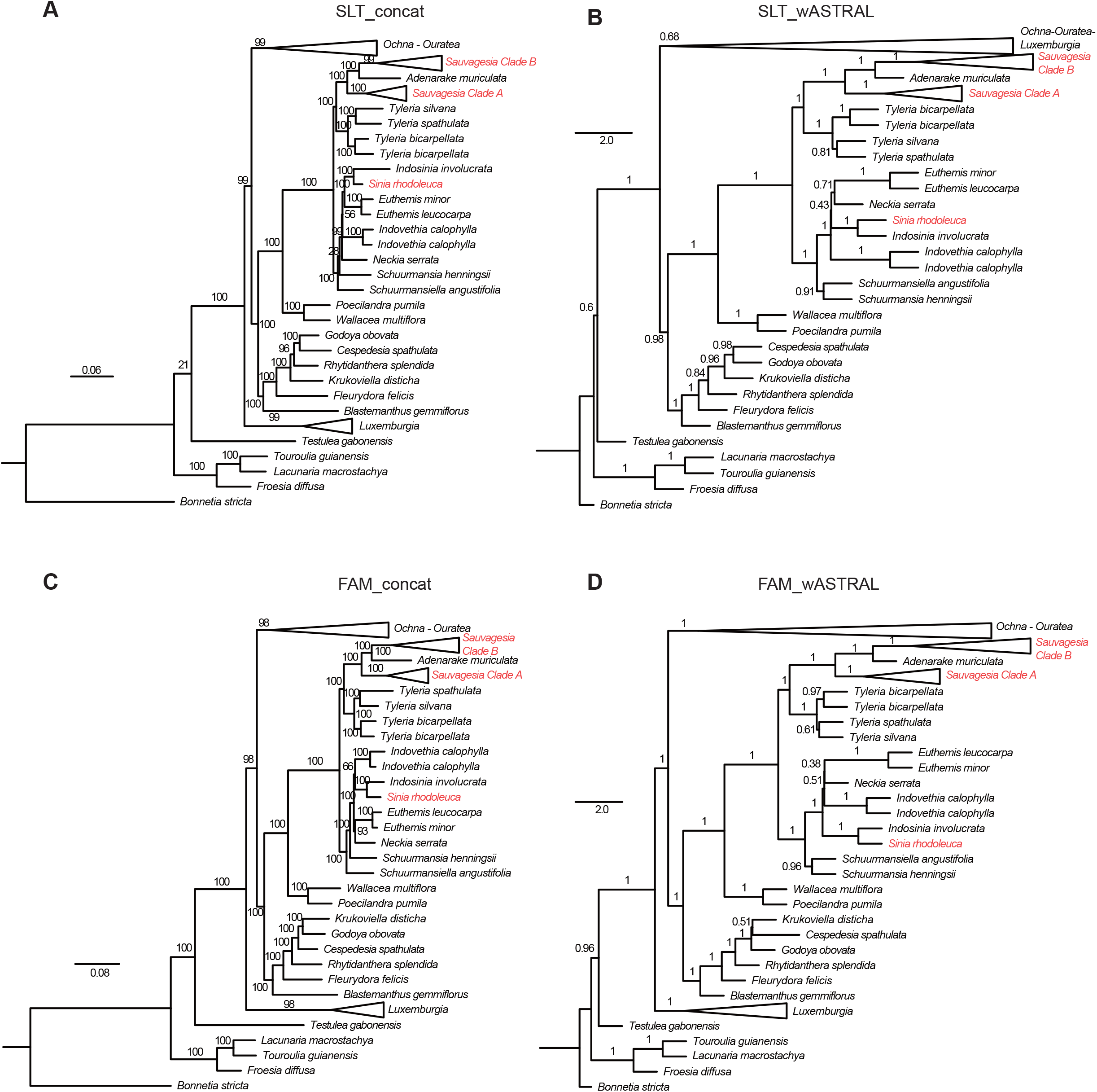
Phylogenetic position of *Sinia rhodoleuca* in tribe Sauvagesieae based on nuclear datasets. SLT, dataset that includes only species from tribes Sauvagesieae, Luxemburgieae, and Testuleeae; FAM, dataset that maximizes taxon sampling across Ochnaceae; concat, tree inference using concatenation method (RAxML); wASTRAL, tree inference using coalescent method (weighted ASTRAL). Number above branch indicates bootstrap value (RAxML) and local posterior probability (weighted ASTRAL)

The plastid phylogeny also supported the separation of *Sinia rhodoleuca* from the sampled *Sauvagesia* species with maximal support (bootstrap = 100%; Fig. 2).

**Figure 2.**
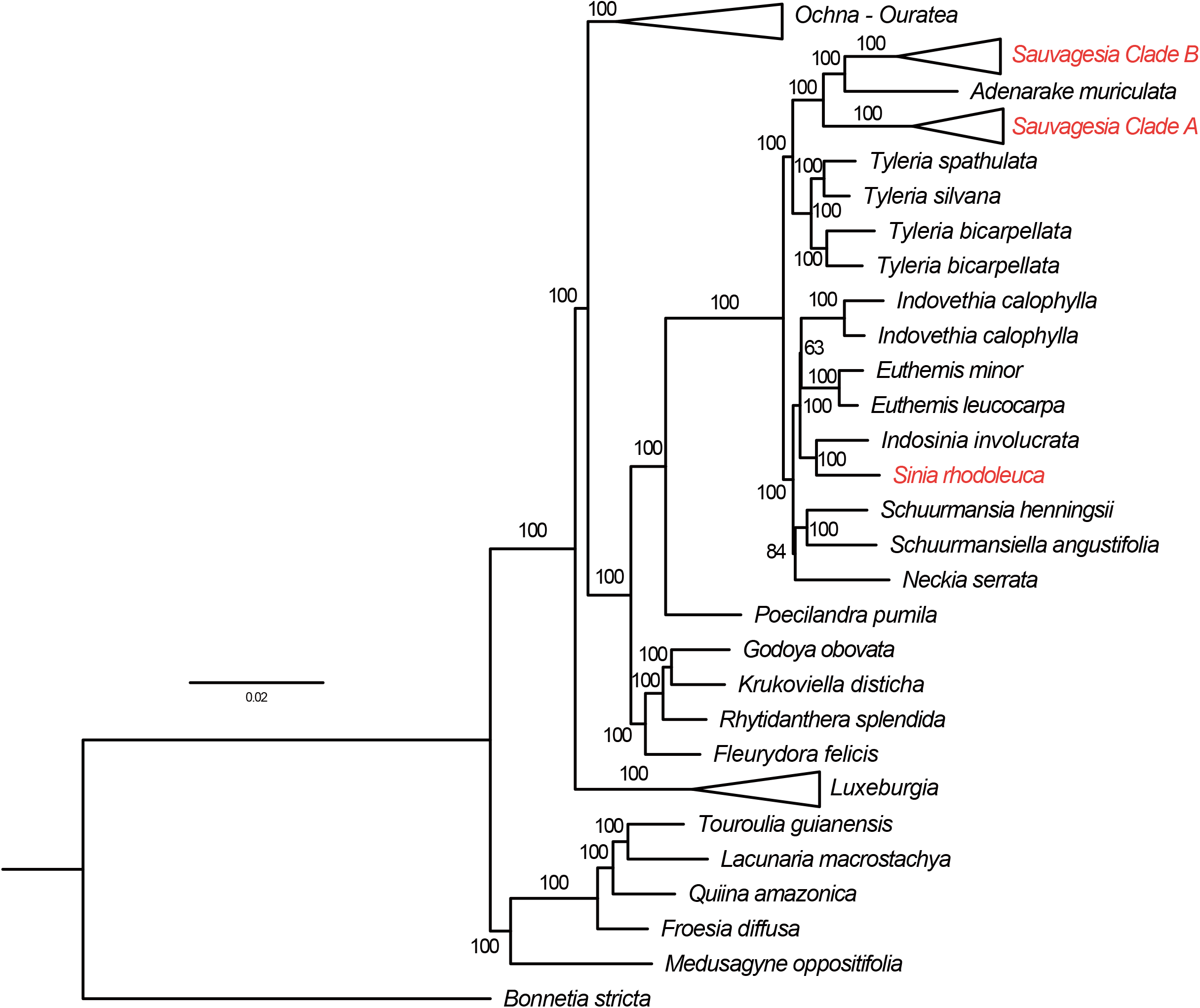
Phylogenetic position of *Sinia rhodoleuca* in tribe Sauvagesieae based on pastid dataset. Number above branch indicates bootstrap value (RAxML)

## Discussion

Our results recovered *Sinia rhodoleuca* as sister to *Indosinia* and revealed that it is distantly related to the Neotropical species of *Sauvagesia*, rendering the currently circumscribed *Sauvagesia* non-monophyletic. This topology presents two alternative taxonomic solutions: either (1) expand *Sauvagesia* to include *Sinia rhodoleuca, Adenarake, Tyleria, Indosinia, Euthemis, Indovethia, Neckia, Schuurmansia*, and *Schuurmansiella*, or (2) reinstate *Sinia* as a distinct genus while retaining the other currently recognized genera. Although broader generic circumscriptions have sometimes been advocated to reduce taxonomic fragmentation (Humphreys and Linder 2009), we argue that reinstating *Sinia* as a distinct genus provides a more informative and morphologically coherent classification.

This treatment is also supported by several morphological similarities between *Sinia* and *Indosinia*. Morphologically, the two genera share lacerate stipules, whereas *Sauvagesia* is characterized by pectinate stipules. Both *Sinia* and *Indosinia* also have numerous, closely parallel secondary veins. Although staminodes are widespread and structurally diverse throughout Sauvagesieae, both genera possess prominent petaloid staminodes that surround the fertile stamens. In *Sinia*, the inner petaloid whorl comprises 10 petaloid staminodes, differentiated into five longer, 3-nerved members and five shorter, 1-nerved members. This arrangement may be comparable to the 10 petaloid staminodes of *Indosinia*, although their homology remains uncertain.

These similarities provide morphological support for the sister relationship between *Sinia* and *Indosinia*, although their phylogenetic significance varies among characters. Lacerate stipules represent a relatively discrete shared feature, whereas closely parallel secondary veins are more widely distributed across Sauvagesieae and may therefore not constitute a synapomorphy of the two genera. Likewise, the elaborate staminode structures of *Sinia* and *Indosinia* are potentially informative, but their evolutionary homology remains uncertain given the considerable diversity and apparent homoplasy of staminode morphology within Sauvagesieae. Despite their close relationship, *Sinia* and *Indosinia* differ in several floral and gynoecial characters, including petal aestivation, anther dehiscence, carpellary number, placentation, and staminode organization. These morphological differences, together with their distinct phylogenetic position, support the recognition of *Sinia* and *Indosinia* as separate genera.

### Taxonomy treatment

#### *Sinia* Diels, reinstated

*Sinia* Diels, Notizbl. Bot. Gart. Berlin-Dahlem 10: 888. 1930.

##### Type

*Sinia rhodoleuca* Diels.

##### Diagnosis

*Sinia* differs from *Indosinia* in having imbricate petals, two whorls of staminodes, longitudinally dehiscent anthers, a 3-carpellate ovary, and parietal placentation.

##### Distribution

Guangdong and Guangxi, China, and northern Vietnam.

***Sinia rhodoleuca*** Diels, Notizbl. Bot. Gart. Berlin-Dahlem 10: 889. 1930 ≡ *Sauvagesia rhodoleuca* (Diels) M.C.E.Amaral, Novon 16: 2. 2006 – **Lectotype** (designated by Amaral, 2006): China. Guangxi (Kwangsi), Yaoshan, Lohsiang, 600–1000 m, 4 May 1929, Sin 8197 (SYS, not seen; photograph at P); **isolectotype**: K.

##### Description

Small erect shrubs, ca. 1 m tall. Stems single or forked near apex, dark purple, striate, glabrous. Stipules (2–)3–5 mm long, margin ciliate, persistent. Leaves alternate; petioles 3–5 mm long, adaxially sulcate, leaving prominent scars on older stems; leaf blades narrowly lanceolate or narrowly elliptic, 7–15 × 1.5–3 cm, papery, both surfaces glabrous, base narrowly cuneate, margin with dense, unequal glandular teeth, apex acute to acuminate; midvein prominent on both surfaces; secondary veins numerous, ± parallel; veinlets conspicuous on both surfaces. Inflorescences paniculate, terminal, 6–10 cm long; peduncles 3–4 cm long. Sepals light green, ovate to lanceolate, 3–4 mm long, margins ciliate-glandular. Petals white or pink, broadly elliptic, 4–6.5 mm long, slightly concave. Stamens 2.5–3.5 mm long; filaments short; anthers sagittate, ca. 2 mm long, longitudinally dehiscent. Staminodes persistent, white, in two whorls, basally slightly connate with the fertile stamens; outer whorl numerous, broadly spatulate, ca. 1 mm long; inner whorl 10, petaloid, oblong, 4–5 mm long, 5 longer and 3-nerved alternating with 5 shorter and 1-nerved. Ovary 3-carpellate, ovoid, 1.5–3 mm long; style terete. Capsule ovoid, ca. 5 mm long, 3-valvate. Seeds ca. 1.5 mm long; testa dark red.

## Acknowledgement

The authors thank Yu-Ying Zhou (South China Botanical Garden) for her kind help in DNA extraction.

## Data availability

Raw sequencing data and plastid genome of *Sinia rhodoleuca* has been deposited and released in National Center for Biotechnology Information (NCBI) Sequence Read Archive (SRA accession number: SRR36996081) and GenBank (accession number: PX933086), respectively.

## Author information

### Authors and Affiliations

**Development Center of Applied Ecology and Ecological Engineering in Guangdong Universities, College of Biology and Food Engineering, Guangdong University of Education, 510303, Guangzhou, China**

Tian-Wen Xiao

**South China National Botanical Garden, Guangzhou 510650, China**

Xue-Jun Ge

**State Key Laboratory of Plant Diversity and Specialty Crops, South China Botanical Garden, Chinese Academy of Sciences, Guangzhou 510650, China**

Xue-Jun Ge

### Contributions

T-W.X. and X-J G. conceived and designed the research; T-W.X. analyzed the data and wrote the original draft; X-J G. reviewed and edited the manuscript; T-W.X. acquired the funding. All authors read and approved the manuscript.

