## Supplementary material for "Phylogenomics supports the reinstatement of *Sinia* (Ochnaceae)": Table S1 and Figure S1

**Table S1** Accession numbers of raw sequencing data generated by previous study.

**Figure S1** Circular map of *Sinia rhodoleuca* plastome. Different functional genes are color coded. LSC, large single-copy region; SSC, small single-copy region; IRa and IRb, two inverted repeats. GC content is indicated by darker gray in the inner circle.
