## Supplementary figures and images for "Phylogenomics supports the reinstatement of *Sinia* (Ochnaceae)"

### Figure S1

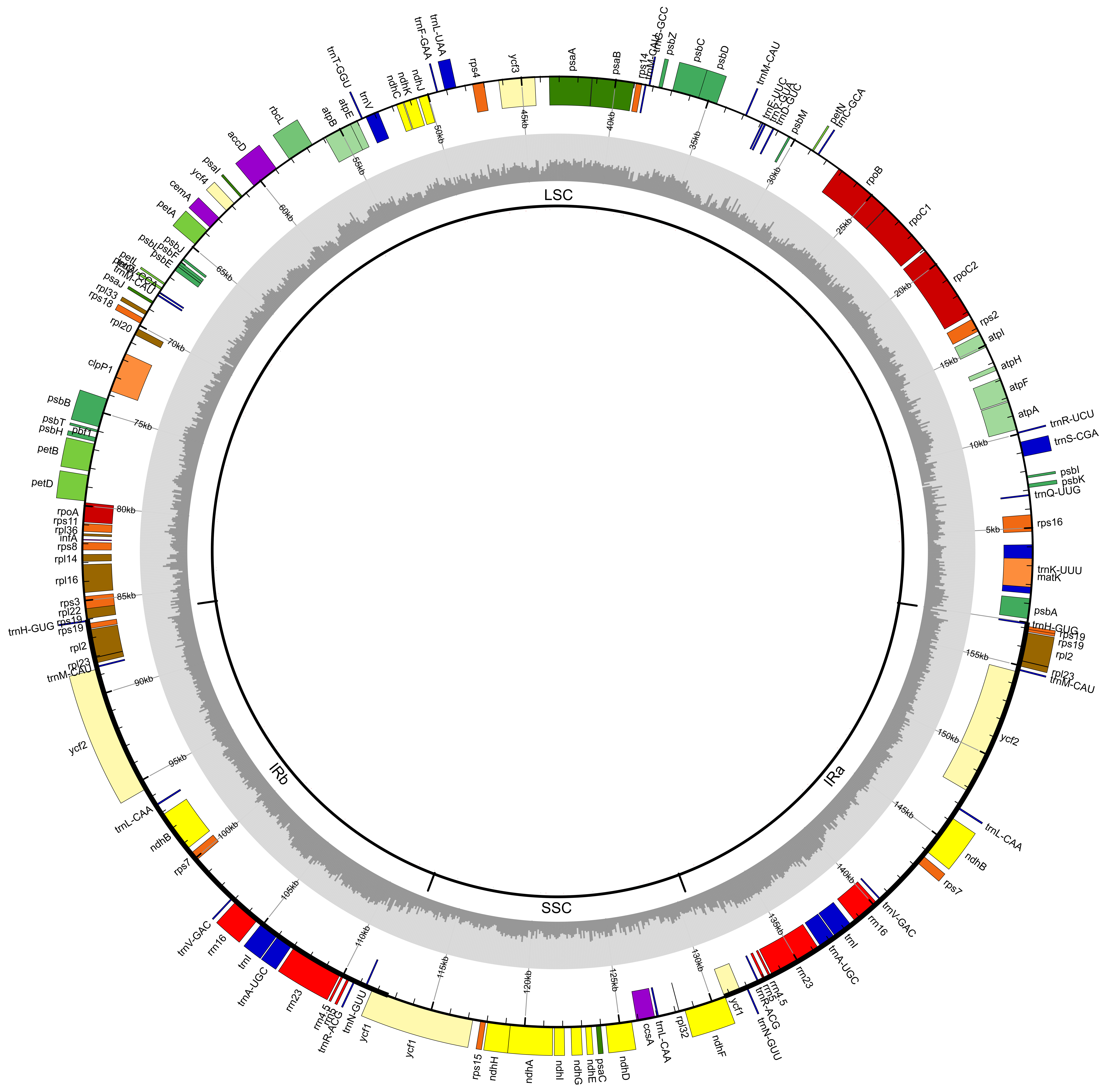
